# Actin and an unconventional myosin motor, TgMyoF control the organization and dynamics of the endomembrane network in *Toxoplasma gondii*

**DOI:** 10.1101/2020.07.15.203950

**Authors:** Romain Carmeille, Aoife T. Heaslip

## Abstract

*Toxoplasma gondii* is an obligate intracellular parasite that relies on three distinct secretory organelles, the micronemes, rhoptries and dense granules, for parasite survival and disease pathogenesis. Secretory proteins destined for these organelles are synthesized in the endoplasmic reticulum (ER) and sequentially trafficked through a highly polarized endomembrane network that consists of the Golgi and multiple post-Golgi compartments. Currently, little is known about how the parasite cytoskeleton controls the positioning of the organelles in this pathway, or how vesicular cargo is trafficked between organelles. Here we show that F-actin and an unconventional myosin motor, TgMyoF, control the dynamics and organization of the organelles in the secretory pathway, specifically ER tubule movement, apical positioning of the Golgi and post-Golgi compartments, apical positioning of the rhoptries and finally, the directed transport of Rab6-positive and Rop1-positive vesicles. Thus, this study identifies TgMyoF and actin as the key cytoskeletal components that organize the endomembrane system in *T. gondii*.

**Author Summary:** Endomembrane trafficking is a vital cellular process in all eukaryotic cells. In most cases the molecular motors myosin, kinesin and dynein transport cargo including vesicles, organelles and transcripts along actin and microtubule filaments in a manner analogous to a train moving on its tracks. For the unicellular eukaryote *Toxoplasma gondii*, the accurate trafficking of proteins through the endomembrane system is vital for parasite survival and pathogenicity. However, the mechanisms of cargo transport in this parasite are poorly understood. In this study, we fluorescently labeled multiple endomembrane organelles and imaged their movements using live cell microscopy. We demonstrate that filamentous actin and an unconventional myosin motor named TgMyoF control both the positioning of organelles in this pathway and the movement of transport vesicles throughout the parasite cytosol. This data provides new insight into the mechanisms of cargo transport in this important pathogen and expands are understanding of the biological roles of actin in the intracellular phase of the parasite’s growth cycle.

## Introduction

*Toxoplasma gondii* is a member of the phylum Apicomplexa, which contains over 5000 species of parasites that cause substantial morbidity and mortality worldwide (1,2). *T. gondii* can cause life-threatening disease in immunocompromised individuals and when infection occurs in utero (3–5). Additionally, *T. gondii* is estimated to cause persistent life-long infection in 10-70% of the world’s population depending on geographic location (6).

*T. gondii* is an obligate intracellular parasite, and thus parasite survival and disease pathogenesis rely on the parasite’s lytic cycle involving host cell invasion, parasite replication within a specialized vacuole termed the parasitophorous vacuole (PV), and host cell egress that results in destruction of the infected cells (reviewed by (7)). To complete this lytic cycle, the parasite relies on three specialized secretory organelles, the micronemes, rhoptries, and dense granules. Micronemes are small vesicles that are localized predominately at the parasite’s apical end (8) and are important for parasite motility, attachment and initiating invasion (Reviewed by (9)). Rhoptries are larger club shaped organelles that contain two sub-sets of proteins (rhoptry bulb proteins (ROPs) and rhoptry neck proteins (RONs)) categorized based on their functions and location within the rhoptry (10). After initial attachment to the host cell, RONs contribute to the formation of the moving junction, a ring structure that aids in the propulsion of the parasite into the host cell (11–15). Once invasion is initiated ROPs and dense granule proteins (GRAs) are secreted into the host cell where they control the organization and structure of the PV, and modulate host gene expression and immune response pathways (10,16–18).

Secretory proteins destined for these distinct organelles are synthesized in the endoplasmic reticulum (ER) and must sequentially traverse multiple intermediate compartments within *T. gondii’s* highly polarized endomembrane system before ultimately arriving at their final destination. Newly synthesized proteins are first trafficked to the Golgi which is located adjacent to the nucleus at the parasite’s apical end. Dense granules are formed from post-Golgi vesicles (19,20) while proteins destined for the micronemes and rhoptries are trafficked through one or more post-Golgi compartments (PGCs), although the exact route taken by each secretory protein has not been elucidated and the function of each PGC has not been fully defined (Fig. 1a). Two of these compartments are marked by Rab5a and Rab7 and are referred to as the endosome-like compartments, as these proteins are markers of the early and late endosomes in higher eukaryotes (21). While the function of the Rab7 compartment is not known (22), overexpression of Rab5a results in the mislocalization of rhoptry proteins and a subset of microneme proteins, to the dense granules, indicating a role in protein sorting (23). Another Rab GTPase, Rab6, is thought to localizes to both the Golgi and the dense granules although the function of Rab6 has not been defined (24). Syntaxin 6 (TgSyn6) marks yet another distinct PGC and appears to have a role in retrograde trafficking between the Rab5 and Rab7 compartments and the trans-golgi network (25). A fifth compartment, the plant-like vacuole (VAC) has similarities to the central vacuole in plants. This compartment contains lytic enzymes that are important for proteolytic processing of the micronemes, ion regulation and processing of endocytic material (26–28). Endocytosis has only recently been described in *T. gondii* (29,30). After uptake material is trafficked to the VAC (31,32). Although the function of PGCs have not been fully defined, these compartments mark a point of intersection between biosynthetic and endocytic pathways and are a hub for protein trafficking in the parasite.

**Figure 1.**
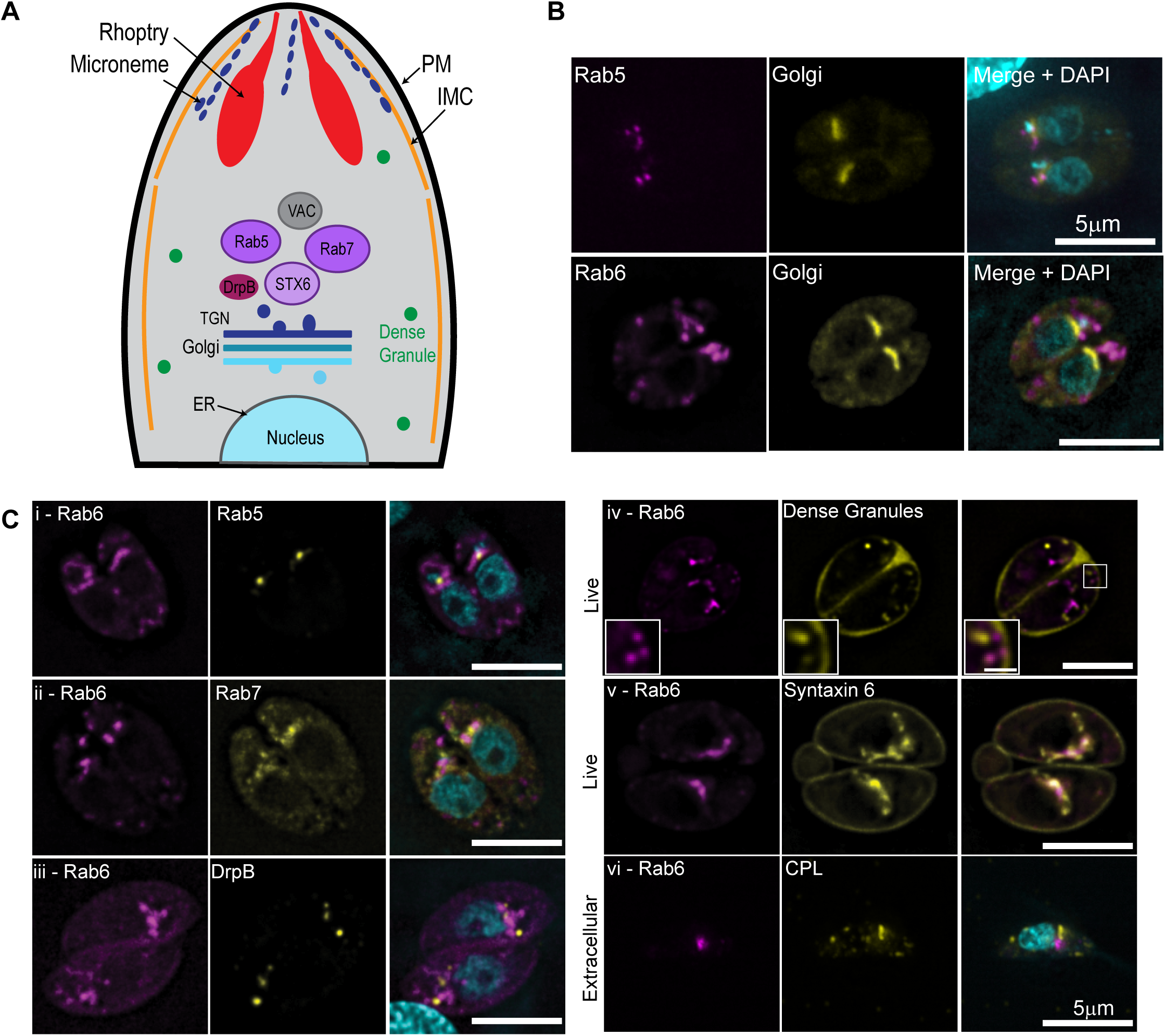
Rab6 and syntaxin 6 co-localize to the same post-Golgi compartment. (A) Schematic diagram of the endomembrane system in *T. gondii*. (B). RH parasites expressing Grasp55-mcherry (Golgi marker; yellow) and NeonFP-Rab5 (magenta; upper panel) or (EmGFP-Rab6 (magenta; lower panel) were fixed and stained with DAPI (cyan). (C) RH parasites expressing AppleFP-Rab6 (magenta; all panels) with markers of the endomembrane system as indicated (yellow). Nuclei were stained with DAPI (cyan). Panels i-iii are deconvolved images of a single focal plane from fixed intracellular parasites. Panels iv-v are a single focal plane from live intracellular parasites expressing AppleFP-Rab6 (magenta) and SAG1ΔGPI-GFP (marker for the dense granules) or Syn6-GFP respectively. Panel vi is a single extracellular parasite expressing AppleFP-Rab6 that fixed and stained with an anti-CPL antibody and DAPI. Scale bar = 5µm. Inset scale bar in panel iv is 1µm.

The location of the PGCs adjacent to the apically positioned Golgi is presumed to optimize the transport of newly synthesized proteins to the parasite’s apical end (22). However, the cytoskeletal factors which control endomembrane positioning have not been identified. In mammalian cells, vesicle transport and organelle positioning is controlled by the microtubule cytoskeleton and associated kinesin and dynein motors (33,34). In contrast, budding yeast uses bundled F-actin filaments as tracks for myosin V-based transport of vesicles, organelles and localizing RNA transcripts (35). *T. gondii* contains 22 highly-stable microtubules (MTs), that are strictly localized at the parasite pellicle (36–38) (Fig. 1a) and do not associate with organelles in the parasite cytosol. Moreover, microtubule depolymerization with oryzalin had no effect on dense granule transport (39) and thus are likely not involved in organelle positioning or vesicle transport between the post-Golgi compartments.

*T. gondii* has a single actin gene (TgAct1) that has 83% similarity with chicken skeletal actin (40). The organization of the actin cytoskeleton in *T. gondii* remained elusive for many years, most likely because of the low propensity TgAct1 to bind phalloidin, the gold standard reagent used to image actin in other cell types (41,42). This inability to visualize F-actin *in vivo*, in addition to biochemical studies which suggested that TgAct1 was incapable of forming long stable filaments *in vitro* led to the idea that short TgAct1 filaments formed only transiently during parasite motility and host cell invasion (reviewed by (43)). However, more recent studies have uncovered new biological roles for TgAct1 including apicoplast inheritance (a non-photosynthetic plastid organelle) (44), dense granule transport (39) and microneme recycling during parasite division (45). Additionally, an unconventional myosin motor, TgMyoF, was shown to be required for apicoplast inheritance and dense granule transport (39,44). With the development of a new reagent, the Actin chromobody™ (ActinCB) (46), actin filaments have now been visualized in both the parasite cytosol (46–48) and in a tubular network which connects individual parasites (46,49). These *in vivo* observations are consistent with recent *in vitro* experiments by Lu and colleagues who used a total internal reflectance (TIRF) microscopy-based approach to demonstrate that Pfact1, actin from the closely related apicomplexan parasite *Plasmodium falciparum*, is capable of transiently forming long filaments up to 30μm in length (50). Collectively, these findings have significantly altered our understandings of both the functions and organization of Apicomplexan actin.

The goal of this study was to investigate the role of actin and TgMyoF in regulating vesicle trafficking and organelle positioning within the endomembrane pathway. Our data demonstrates that both of these proteins are required for the apical positioning and morphology of the Golgi and post-Golgi compartments, ER tubule movement, and transport of Rab6-positive and Rop1-positive vesicles. These results indicate that this acto-myosin system is vital for controlling the organization of the endomembrane system in *T. gondii* and uncovers new biological roles of actin in the intracellular phase of the parasite’s growth cycle.

## Results

### Rab6 colocalizes with syntaxin6 in a post-Golgi compartment

To investigate how the morphology and dynamics of the PGCs is controlled by the *T. gondii* cytoskeleton, we fluorescently labeled the Rab5a and Rab6 PGCs by expressing NeonGreenFP-Rab5a (referred to subsequently as Neon-Rab5a) and EmeraldFP-Rab6 (EmGFP-Rab6) along with Grasp55-mCherryFP, a marker of the cis-Golgi (23,24,51,52). As expected, Neon-Rab5 did not co-localize with Grasp55-mCherryFP and was found in a distinct compartment adjacent to the Golgi (23) (Fig.1B; upper panel). Surprisingly, Rab6 also did not localize to the Golgi as previously reported (24) but also localized to a compartment apical to the Golgi (Fig. 1B; lower panel). To determine if Rab6 colocalizes with markers of the other post-Golgi compartments, we transfected parasites with AppleFP-Rab6 expression construct along with Neon-Rab5, Neon-Rab7, Syn6-GFP, DrpB-GFP and SAG1ΔGPI-GFP, a marker for the dense granules (23,31,53,54) and grew parasites for ∼18 hours before fixation or live cell imaging (Fig. 1C). Additionally, extracellular parasites expressing EmGFP-Rab6 were fixed and stained with an anti-TgCPL antibody, a marker of the VAC (Fig. 1C) (27). Rab6 and Syn6 localize within the same post-Golgi compartment. Syn6 was also localized to the plasma membrane but no peripheral staining of Rab6 was observed (Fig. 1C) (25). No colocalization was observed between Rab6 and the other proteins analyzed (Fig. 1C) including the dense granules which were previously reported to contain Rab6 on their surface (25) (Fig. 1C; panel iv).

Live cell imaging of parasites expressing Syn6-GFP and AppleFP-Rab6 demonstrates that these proteins occupy distinct sub-domains within the same tubular-vesicular compartment (Fig. 2A and Fig. 2B; Video 1). Line scan analysis indicates that Syn6 and Rab6 are both found in the “vesicular” domain of the compartment (Fig. 2B; magenta arrow) while Rab6 positive/Syn6 negative (Rab6(+)/Syn6(-)) tubules extend from this “vesicular” domain (Fig. 2B; white bracket). Rab6(+) vesicles can be seen budding from the tip of the tubular extensions (Fig. 2C). Rab6(+)/Syn6(-) vesicles are distributed throughout the parasite cytosol (Fig. 2D arrowhead; video 1) and exhibited directed, presumably motor-driven, motion.

**Figure 2.**
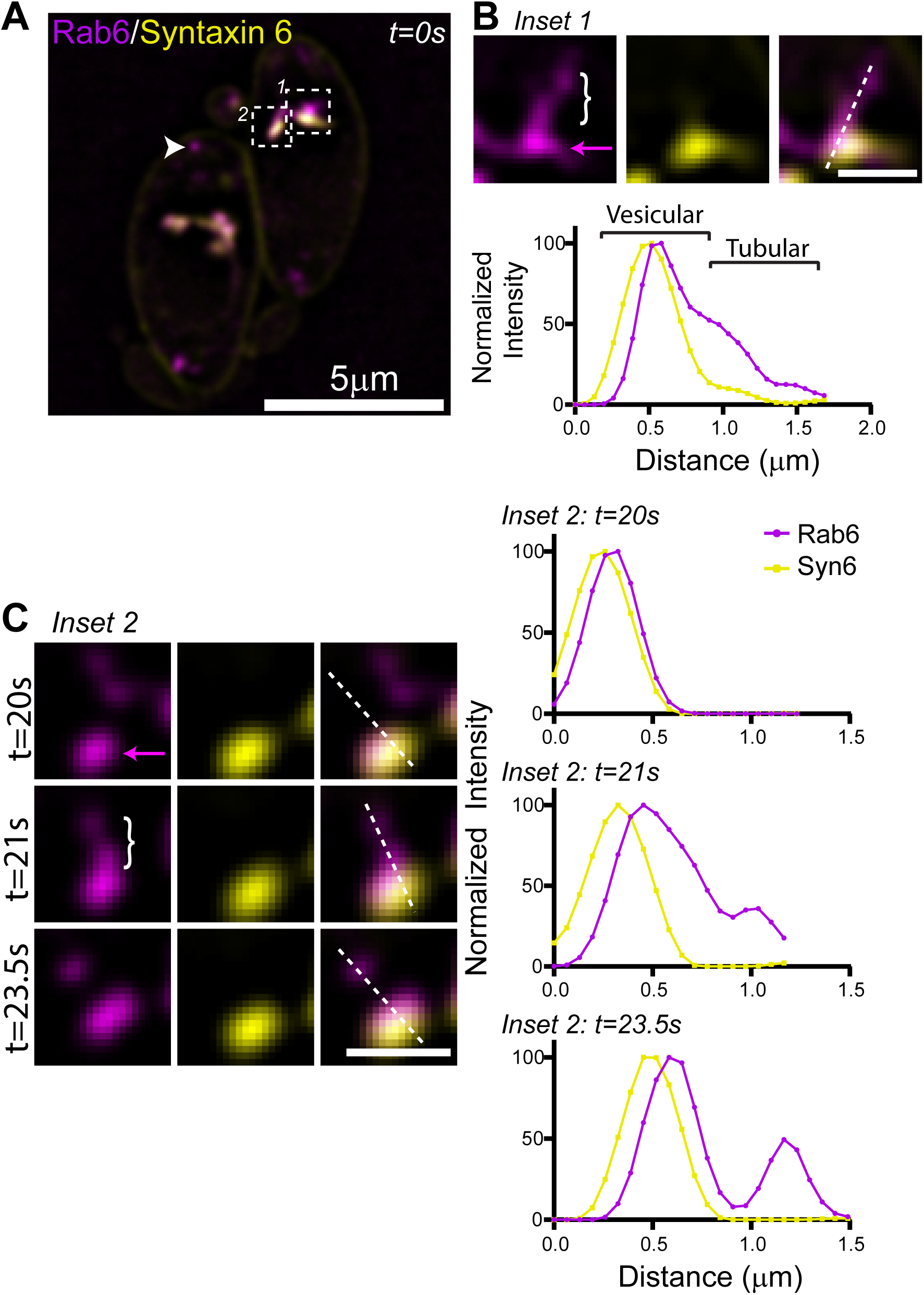
The Rab6 and syntaxin 6 PGC is dynamic. (A) Live imaging of RH parasites expressing AppleFP-Rab6 and Syn6-GFP. Rab6 vesicles indicated with the arrow head. Areas in the white box were used to make inset images in panels (B) and (C). Scale bar = 5µm. (B) Upper panel. Inset 1 from panel (A). Dashed line was used to make line scan in lower panel. Bracket indicates Rab6 positive and Syn6 negative tubule. Magenta arrow indicates vesicular compartment containing both Rab6 and Syn6. Lower panel. Line scan of Rab6 and Syn6. Shift in peaks between Rab6 and Syn6 likely due to sequential image acquisition between the two imaging channels. Scale bar is 1µm. (C) Left panel. Image from Inset 2 from panel (A), taken at times indicated. Bracket indicates Rab6 positive and Syn6 negative tubule. Magenta arrow indicates vesicular compartment containing both Rab6 and Syn6. Dashed lines were used to make line scan in right panel. Right panel. Line scans of Rab6 and Syn6 at times indicated. Scale bar is 1µm.

### Rab6 vesicle transport is dependent on F-actin

To further characterize the dynamics and morphology of the Rab6 compartment, we imaged parasites expressing EmGFP-Rab6 using live cell microscopy with a temporal resolution of 100ms (Fig. 3A; Video 2). The Rab6 compartment is dynamic and undergoes constant rearrangement as indicated by images taken after 5, 10 and 15 seconds of imaging (Fig. 3A, middle panel; Video 2). Rab6(+) vesicles exhibited directed movement throughout the cytosol as illustrated by kymographs (Fig. 3A, right panel). This motion was reminiscent of the directed actin-based motion exhibited by dense granules (39). Tracking the movement of Rab6(+) vesicles revealed velocities and run-lengths of 0.92±0.01μm/s and 1.6±0.03μm respectively (Table S1).

**Figure 3.**
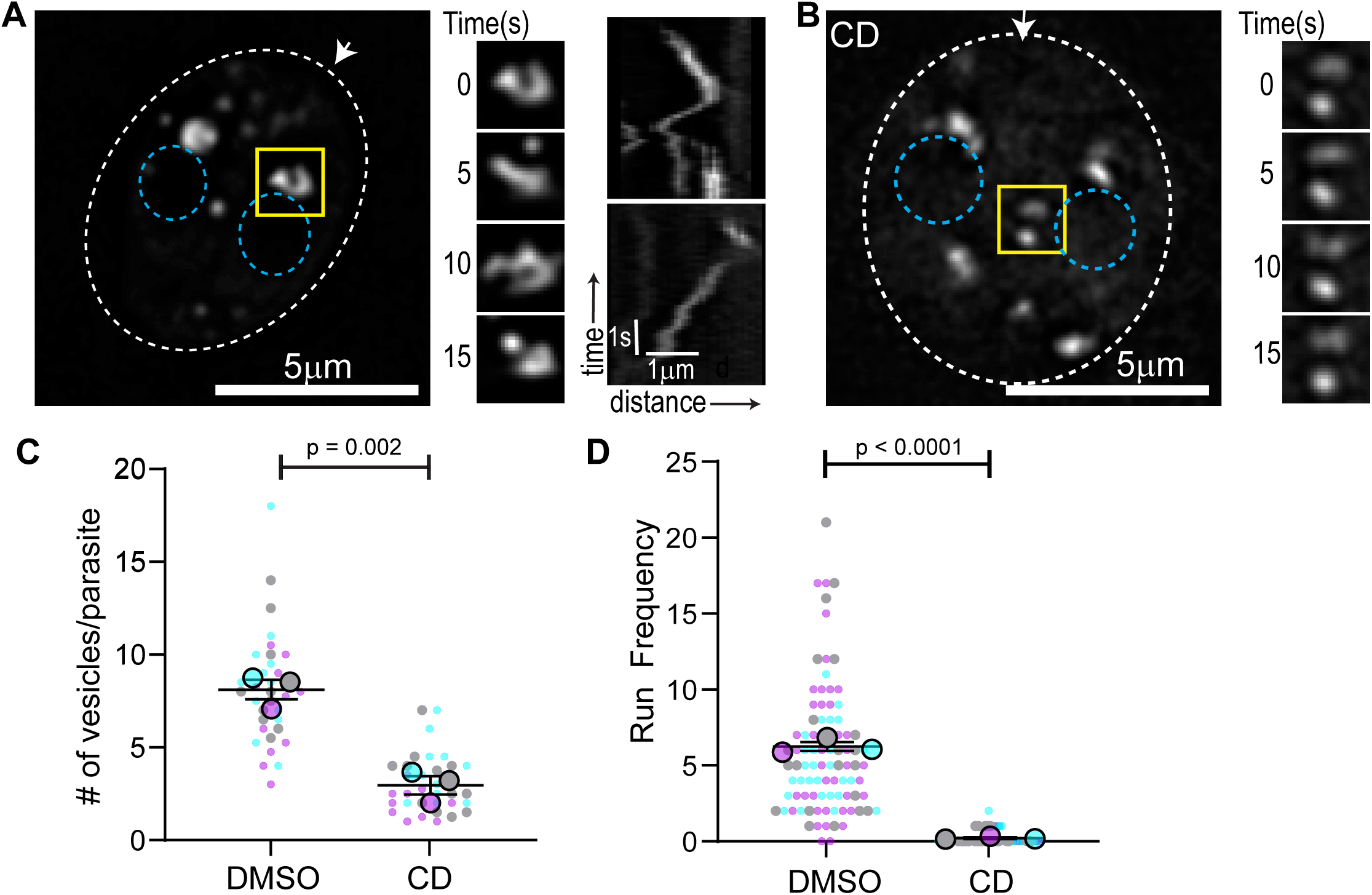
Rab6 vesicle transport is actin dependent. (A-B) *Left panel*. RH parasites expressing EmGFP-Rab6 treated with DMSO (A) or cytochalasin D (CD) (B). Dashed oval indicates the PV surrounding a 2-parasite vacuole. Location of the nucleus is indicated by the blue circles. Area in the yellow box was used to make inset (middle panel). Arrow indicates parasite’s apical end. *Middle panel*. Rab6 compartment dynamics. Images taken at 5 second intervals. *Right panel*. Kymograph depicting Rab6 directed vesicle motion. No directed motion was observed after cytochalasin D treatment. (C) Number of Rab6 vesicles per parasite was quantified in DMSO and CD treated parasites. (D) Number of directed runs/parasite/minute (run frequency) observed in DMSO and CD treated parasites. In (C) and (D) results are from three independent experiments. Mean from each independent experiment is indicated with large circles. Raw data is shown with smaller colored circles. Experiment 1 in magenta, experiment 2 in cyan, experiment 3 in grey. Error bars indicated mean and SEM.

Given the similarities between Rab6(+) vesicle movement and actin dependent dense granule movement (39) we sought to determine if actin was required for Rab6(+) vesicle transport. We treated intracellular parasites expressing EmGFP-Rab6 with cytochalasin D (CD) for 60 minutes to depolymerize F-actin before commencement of imaging (Video 2). We observed two phenotypes associated with the loss of F-actin: First, the main Rab6(+) compartment lost its apical localization and became fragmented and distributed throughout the parasite cytosol (Fig. 3B). Second, the dynamic tubular morphology of the EmGFP-Rab6 compartment is lost and the compartment remained static throughout the 60-second imaging period. Vesicle formation from the Rab6 compartment was perturbed in CD treated parasites (Fig. 3B; right panel). The number of Rab6(+) vesicles in the parasite cytosol decreases from an average of 8±0.5 in control parasites (RH parasites treated with DMSO) to 3±0.3 after CD treatment (Fig. 3C). Similarly, the number of directed runs exhibited by Rab6(+) vesicles decreased from 6±0.45 in control to less than 1 after CD treatment (Fig. 3D).

### Creation of conditional TgMyoF-knockdown parasite line

Since dense granule transport is dependent on F-actin and TgMyoF (39), we investigated if TgMyoF was also required for Rab6(+) vesicle transport or compartment dynamics. We previously used an inducible Cre-LoxP system to create a parasite line deficient in functional TgMyoF, however depletion of TgMyoF protein levels after TgMyoF gene excision took ∼48 hours (39,55). Thus, we created an inducible TgMyoF knockdown (KD) parasite line using the auxin-inducible degradation system where TgMyoF protein levels could be rapidly degraded (56,57) (Fig. 4A). The endogenous TgMyoF gene was C-terminally tagged with an AID-HA epitope to create a TgMyoF-mAID-HA parasite line (referred to subsequently as TgMyoF-AID). PCR of genomic DNA and western blot was used to confirm the accurate integration of this construct (Fig. 4B and Fig. S1). Treatment of TgMyoF-AID parasites with indole-3-acetic acid (IAA) for four hours resulted in depletion of TgMyoF to undetectable levels (Fig. 4B). TgMyoF depletion was independently confirmed by anti-HA immunofluorescence (IF) (Fig. 4C) and TgMyoF-KD parasites exhibited aberrant apicoplast inheritance as demonstrated previously (Fig. S1B) (39,44).

**Figure 4.**
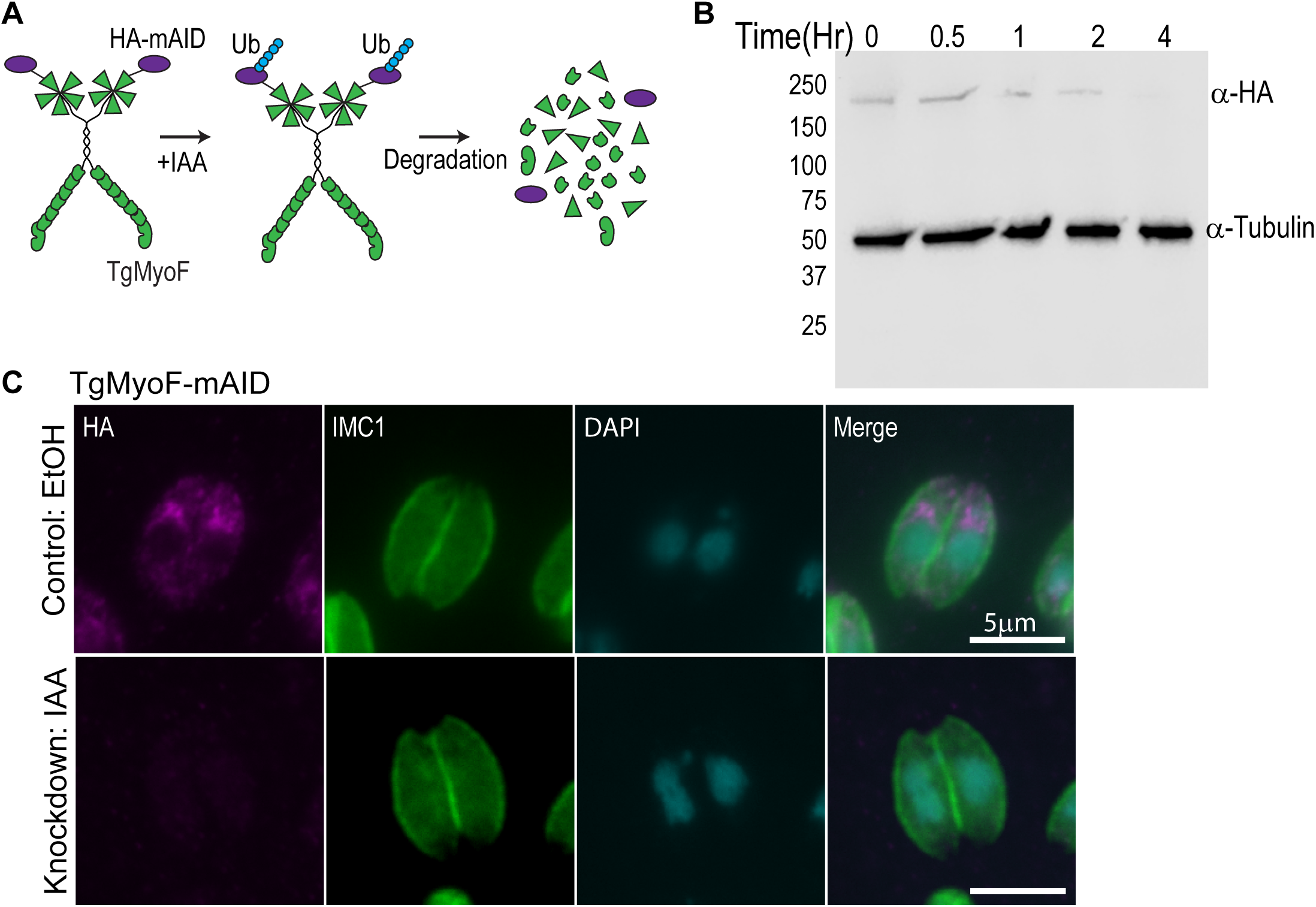
Creation of TgMyoF-KD parasites using the auxin-inducible degradation system. (A) Cartoon of TgMyoF knockdown strategy. (B) Western blot analysis of TgMyoF-AID-HA depletion as a function of IAA treatment time. Tubulin was used as the loading control. (C) Deconvolved epifluorescence images of TgMyoF-AID parasites treated with EtOH or IAA for 15 hours before fixation. Anti-HA immunofluorescence (magenta) was used to assess TgMyoF depletion. Anti-IMC1 antibody (green) and DAPI (cyan) staining used to visualize the parasite periphery and nuclei respectively.

### TgMyoF is required for apical positioning and structural integrity of the PGCs

We were intrigued by the observation that F-actin was required for the apical positioning of the Rab6 compartment. The post-golgi compartments have well-defined positions adjacent to the Golgi, yet we have no mechanistic insight into how the position of these organelles is maintained. To determine if TgMyoF is required for the apical positioning of the PGCs, we ectopically expressed markers for these compartments, specifically EmGFP-Rab6, Neon-Rab5a, Neon-Rab7, Syn6-GFP and DrpB-GFP in TgMyoF-AID parasites treated with ethanol (EtOH; control) or IAA (to deplete TgMyoF) for 18 hours before fixation. As expected in control parasites, these compartments were positioned at the apical end of the parasite (Fig. 5A left panels). After TgMyoF depletion however, the Rab5a, Rab6, DrpB and Syn6 compartments became fragmented and were found throughout the cytosol (Fig. 5A; right panels). For each protein, we quantified the number of compartments per parasite and found a statistically significant increase after TgMyoF depletion in each case (Fig. 5B). In the case of Rab7, this protein had a diffuse localization in the cytosol after TgMyoF depletion (Fig. 5A). Since Rab6 and Syn6 are localized to the same compartment in control parasites, we sought to determine if these proteins remained colocalized after compartment fragmentation. In TgMyoF deficient parasites expressing AppleFP-Rab6 and Syn6-GFP, we found that the co-localization between these proteins remained after compartment fragmentation (Fig. 5C). In addition, 70% of parasites also contained Rab6+/Syn6-vesicles (Fig. 5C, lower panel, white arrows).

**Figure 5.**
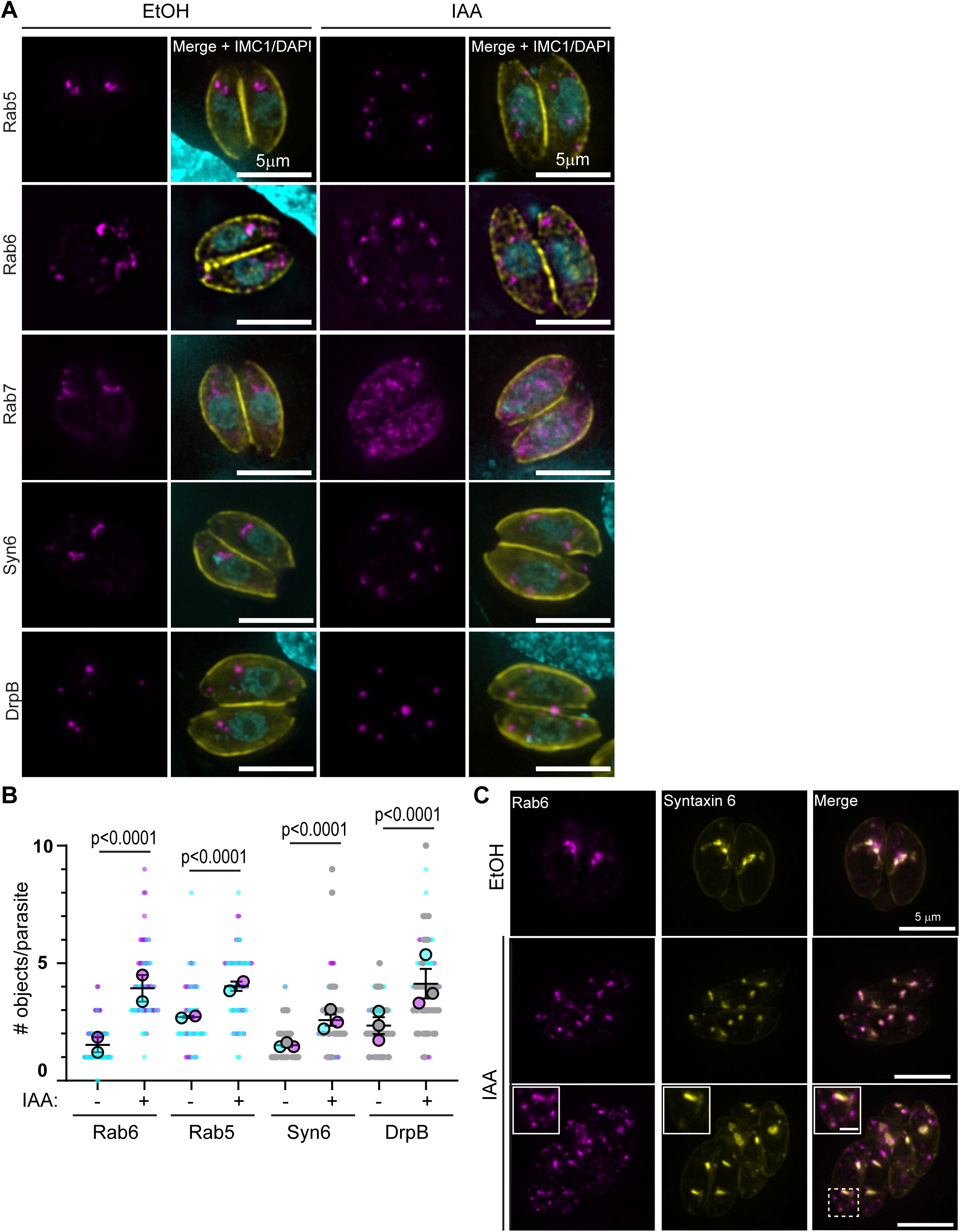
TgMyoF knockdown results in fragmentation and dispersion of post-Golgi compartments. (A) TgMyoF-AID parasites expressing Neon-Rab5, EmGFP-Rab6, Neon-Rab7, GFP-Syn6 or GFP-DrpB (magenta) were treated with EtOH (left panels) or IAA (right panels) for 18 hours before fixation. Parasites were stained with anti-IMC1 to highlight the parasite periphery (yellow) and DAPI to stain DNA (cyan). Scale bar is 5μm. (B) For each post-Golgi compartment, the number of objects was quantified in control and TgMyoF knockdown parasites. Combined results from two or three independent experiments. Mean from each independent experiment is indicated with large circles. Raw data is shown with smaller colored circles. Experiment 1 in magenta, experiment 2 in cyan, experiment 3 in grey. Error bars indicated mean and SEM. (C) TgMyoF-AID parasites expressing Syn6-GFP (yellow) and AppleFP-Rab6 (magenta) treated with EtOH (top row) or IAA (middle and bottom rows). *Bottom row*. Inset is magnification of area in the dashed box. Inset scale bar is 1µm.

### TgMyoF plays a role in Rab6 vesicle transport

To assess the role of TgMyoF in the Rab6 compartment and Rab6(+) vesicle dynamics, TgMyoF-AID parasites expressing EmGFP-Rab6 were treated with either EtOH or IAA for 18 hours before live cell imaging. Similar to what was observed after actin depolymerization, loss of TgMyoF resulted in fragmentation of the Rab6 compartment (Fig. 5A and 5B; left panels) and loss of this compartments dynamic reorganization. (Fig. 6A and 6B middle panel). There was also a decrease in the number of Rab6(+) vesicles exhibiting directed motion from 7.6±0.7 in the control to 3±0.45 in TgMyoF-KD parasites (Fig. 6A and 6B right panels; Fig. 6C) (Video 3). Although the number of directed runs was significantly reduced, vesicle velocities were the same in the absence of TgMyoF compared with controls (Fig. 6D) (Table S1).

**Figure 6.**
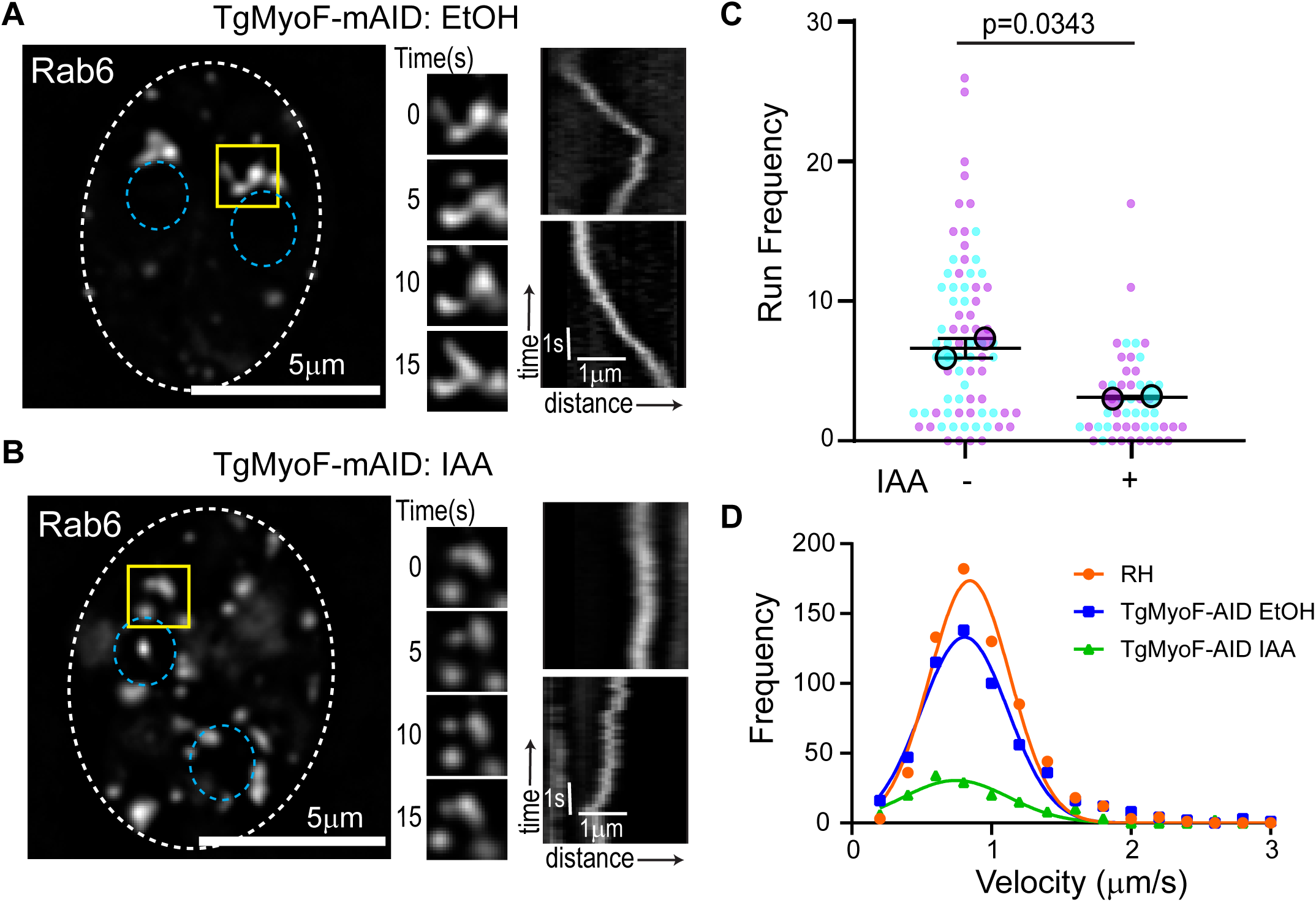
TgMyoF is required for Rab6 vesicle transport. (A-B) *Left panel:* TgMyoF-AID parasites expressing EmGFP-Rab6 treated with EtOH (A) or IAA (B) for 18 hours before imaging. Dashed oval indicates the PV surrounding a 2-parasite vacuole. Location of the nucleus is indicated by the blue circles. Area in the yellow box was used to make inset (middle panel). *Middle panel:* Images of the Rab6 compartment taken at 5 second intervals. *Right panel:* Kymograph depicting Rab6 vesicle motion. (C) Run frequency (# of directed runs/parasite/minute) of Rab6 vesicles in control and TgMyoF depleted parasites. Combined results from two independent experiments. Mean from each independent experiment is indicated with large circles. Raw data is shown with smaller colored circles. Experiment 1 in magenta, experiment 2 in cyan. Error bars indicated mean and SEM. (D) Frequency distribution of Rab6 vesicle velocities in RH parasites (orange) and in TgMyoF-AID parasites treated with EtOH (blue) or IAA (green) for 18 hours.

These data combined with published work demonstrate that TgMyoF and actin are required for dense granule and Rab6+ vesicle transport (39), apical positioning of the post-golgi compartments, and inheritance of the apicoplast (44). All of these organelles are part of the endomembrane network in *T. gondii*. Therefore, we wanted to determine if other organelles in this pathway, namely the ER, the Golgi, the micronemes and the rhoptries, relied on this acto-myosin system for their dynamics and/or morphology.

### TgMyoF-knockdown affects movement of ER tubules

The ER is a large membrane bound organelle that has three distinct functional domains, the nuclear envelope and peripheral tubules and peripheral cisternae which form an extensive and continuous network in the parasite cytosol (52,58). Live cell imaging of parasites with a fluorescently labeled ER, achieved by expression eGFP-SAG1ΔGPI-HDEL (53) (referred to subsequently as GFP-HDEL) reveals that the ER tubules are highly dynamic and undergo continuous reorganization (Video 4). ER tubule rearrangements are clearly evident when the first frame of the movie was overlaid with images taken after 5, 10 and 15 seconds of imaging (Fig 7A; right panel). ER tubule motility has been described previously in mammalian cells (59). In this case, tubule dynamics are dependent on the microtubule cytoskeleton (60–62). To determine if TgMyoF plays a role in ER dynamics in *T. gondii*, TgMyoF-AID parasites expressing GFP-HDEL were imaged after TgMyoF depletion. Loss of TgMyoF did not result in reorganization or collapse of the peripheral ER, however ER tubule motility is decreased as illustrated with time overlay images (Fig. 7B; right panel) (Video 4).

**Figure 7.**
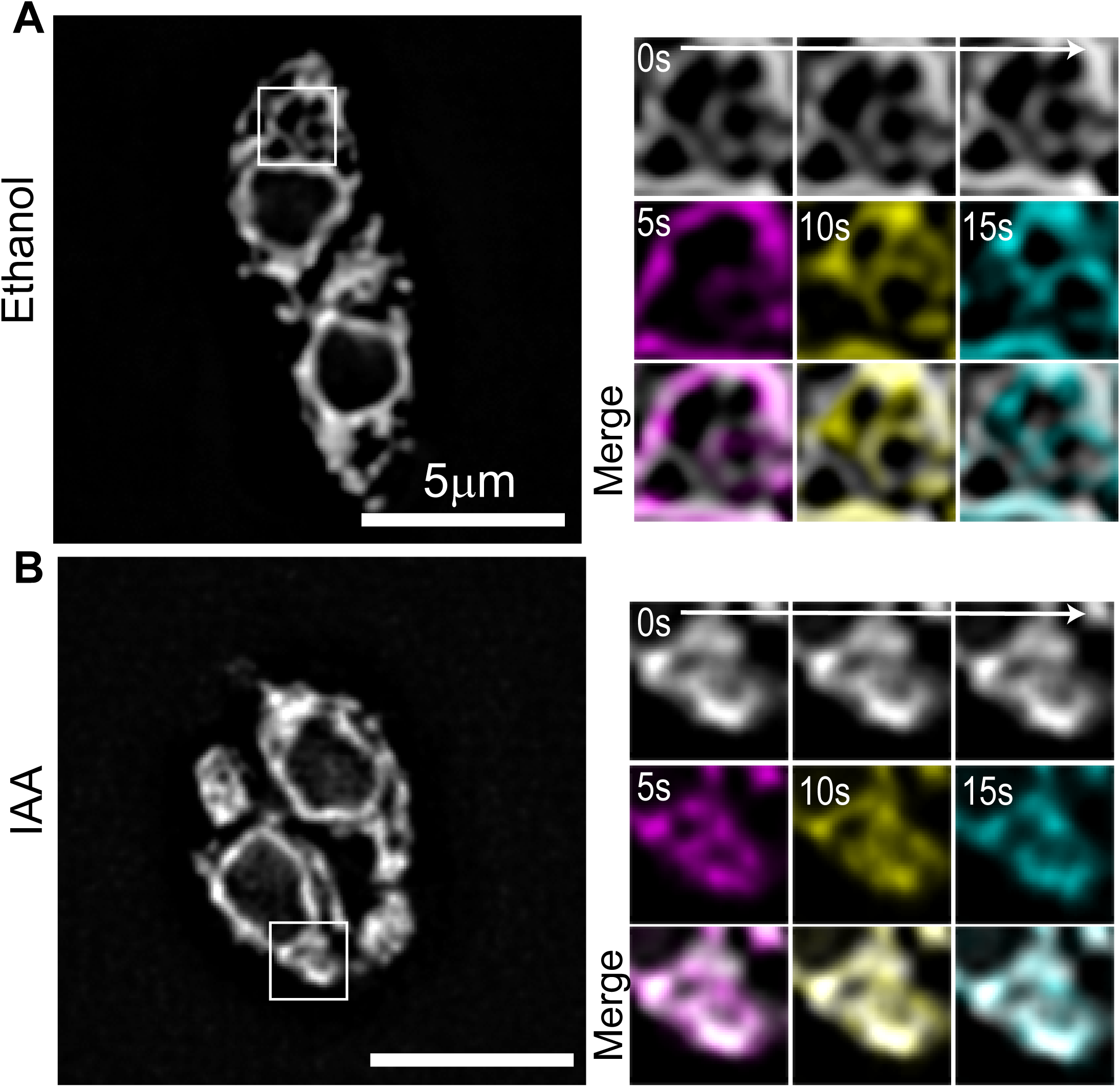
TgMyoF is required for ER tubule movement. (A&B) *Left panel*: TgMyoF-AID parasites treated with EtOH (A) or IAA (B) where the ER is fluorescently labeled with eGFP. *Right panel:* magnification of area in the white box. First image of each movie (grey) was overlaid with images taken after 5s (magenta), 10s (yellow) and 15s (cyan) of imaging.

### Actin depolymerization and TgMyoF depletion results in Golgi fragmentation

Next, we investigated if loss of TgMyoF or actin depolymerization affected Golgi morphology. The cis-golgi was fluorescently labeled by expressing Grasp55-mCherryFP in TgMyoF-AID parasites. Parasites were treated in EtOH or IAA for 15 hours to deplete TgMyoF. In the presence of TgMyoF, 80% of parasites contained a single Golgi localized at the apical end of the nucleus while 20% of parasites were undergoing cell division and contain two Golgi per parasite as expected (Fig. 8A, 8B & 8D). After TgMyoF knockdown, only 25% of parasites contained a single Golgi, 52% contained two Golgi and 21% contained three Golgi (Fig. 8B and 8D). Similarly, actin depolymerization with cytochalasin D also resulted in an increased number of Golgi per parasite (Fig. 8C and 8D). While the majority of Golgi remain closely associated with the nucleus after TgMyoF-knockdown or CD treatment, the apical positioning of the Golgi was lost. After TgMyoF depletion or actin depolymerization, 52% and 40% of parasites respectively contained Golgi in both the apical and basal ends of the parasites compared to just 5% of control parasites (Fig 8E).

**Figure 8.**
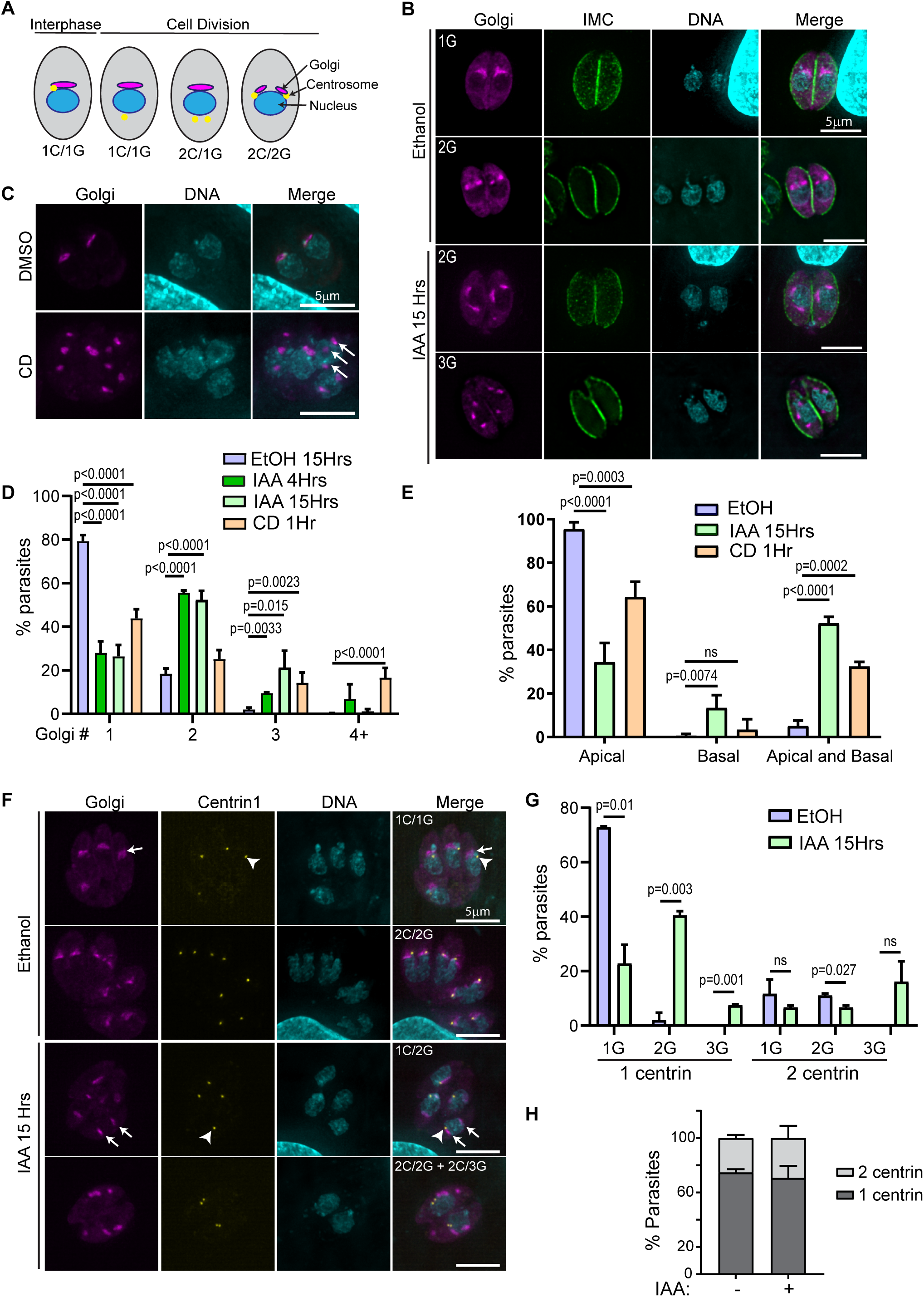
TgMyoF and actin control Golgi morphology. (A) Diagram of centrosome and golgi division. (B) TgMyoF-AID parasites expressing mCherry-Grasp55 (a marker of the cis-Golgi; magenta) were fixed and stained with anti-IMC1 (green) and DAPI (cyan) after 15 hours of EtOH or IAA treatment. (C) TgMyoF-AID parasites expressing mCherry-Grasp55 were treated with DMSO or CD for 60 minutes before fixation and DAPI staining. White arrows indicate Golgi fragments. (D) Quantification of # of Golgi/parasite in TgMyoF-AID parasites treated with EtOH for 15 hours (blue), IAA for 4 hours (dark green), IAA for 15 hours (light green) or CD for 60 minutes (orange). Results are combined data from at least 2 independent experiments. N>50 parasites/experiment. (E) Quantification of Golgi localization in apical only, basal only or both the apical and basal ends of the parasite. Results are combined data from 2 independent experiments. N>50 parasites/experiment. (F) TgMyoF-AID parasites expressing mCherry-Grasp55 (magenta) and centrin 1-GFP (as a marker of the centrosome; yellow) were treated with EtOH and IAA for 15 hours before fixation and DAPI staining. Arrowhead indicates the centrosome. Arrows indicate the Golgi. (G) Quantification of centrosome and Golgi number in control and TgMyoF depleted parasites. Combined results from 2 independent experiments. N=77 parasites for control, N=93 parasites for IAA treated. (H) Centrosome number per TgMyoF-AID parasite treated with EtOH or IAA. Combined results from 2 independent experiments. N=77 parasites for control, N=93 parasites for IAA treated.

Since the Golgi in *T. gondii* divides by binary fission during cell division (51,63), the increased number of Golgi observed after the loss of F-actin and TgMyoF could be due to uncoupling of the Golgi division cycle from the cell cycle. To determine if this was the case, we first determined the number of Golgi per parasite after 4 hours of IAA treatment, the time at which TgMyoF is completely depleted and a length of time shorter than one parasite division cycle. After 4 hours and 15 hours of IAA treatment, the number of Golgi per parasite is indistinguishable indicating that Golgi fragmentation occurs quickly upon TgMyoF depletion but does not continue to fragment with extended IAA treatment times (Fig 8D). Next, we investigated if loss of TgMyoF affected the number of centrosomes per parasite as it had previously been demonstrated that centrosome duplication and Golgi fission are the first events to take place at the start of parasite division (52) (Fig. 8A). TgMyoF-AID parasites were transfected with Grasp55-mCherry and centrin1-GFP (64) to label the Golgi and centrosomes respectively, and then treated with EtOH or IAA for 15 hours. In control parasites, 73% of parasites contained one Golgi and one centrin (1G/1C) while the remaining ∼30% of parasites were at various stages of division and contained either one Golgi and twp centrin (1G/2C) or two Golgi and two centrin (2G/2C) (Fig. 7F and 7G). In contrast, only 20% of TgMyoF-KD parasites contained one Golgi and one centrin (1G/1C), while 41% contained two Golgi and one centrin (2G/1C), compared to just 3% of controls. 7% and 16% of IAA treated parasites contained one centrin and three Golgi (1C/3G) or two centrin and three Golgi (2C/3G) respectively, which were phenotypes that were never observed in the control parasites (Fig. 8F and 8G). The number of centrosomes per parasite remained unchanged in TgMyoF-KD parasites compared to controls (Fig. 8H). Thus, we conclude that TgMyoF and actin are important for controlling both Golgi number and apical positioning and these effects on Golgi morphology are independent of the parasite’s cell division cycle.

### TgMyoF depletion affects apical positioning of the rhoptries and Rop1 vesicle movement

It has been demonstrated previously that loss of TgMyoF results in accumulation of intact micronemes and rhoptry proteins in the parasite’s residual body (44). In our independently generated TgMyoF conditional knockdown parasite line, we observe a similar phenotype. After 15 hours of IAA treatment, 73% of TgMyoF deficient vacuoles contained rhoptries (visualized by expression of Rop1-NeonFP) in the residual body compared to just 14% of controls. While 67% of TgMyoF deficient vacuoles exhibited accumulation of micronemes in the residual body (visualized using an anti-AMA1 antibody) compared to 5% of controls (Fig. 9A and 9B). Despite the accumulation of these organelles in the residual body, the apical positioning of the micronemes was not affected in TgMyoF-knockdown parasites when assessed by IFA (Fig. 9A). In contrast, there appeared to be an increased Rop1 fluorescence throughout the parasite cytosol (Fig. 9A; magenta arrow). To further investigate the effects of TgMyoF depletion on rhoptry dynamics, we expressed Rop1-NeonFP in TgMyoF-AID parasites treated for 18 hours with either EtOH or IAA and imaged the parasites using live cell microscopy. In control parasite’s, the rhoptries were localized as expected at the apical end. The rhoptries were surprisingly dynamic, and like the Rab6 compartment, were constantly rearranged (Fig. 9C, inset; Video 5). In addition, Rop1 vesicles were observed throughout the parasite and exhibited directed, motor-driven motion (Video 6). Upon TgMyoF knockdown we observed a large decrease in the number of directed runs exhibited by Rop1-NeonFP vesicles from 11±1.1 in control parasites to 1.4±0.2 after IAA treatment, even though the total number of Rop1 vesicles was not statistically different between control and TgMyoF depleted parasites (6.25±0.4 and 9.1±0.6 in EtOH and IAA treated cells respectively) (Fig. 9E and 9F). To further investigate the effect of TgMyoF knockdown on the apical positioning of the rhoptries, we compared Rop1-NeonFP fluorescence intensity at the apical and basal ends in control and TgMyoF depleted parasites. In control parasites the apical:basal ratio was 5.8±0.7, indicating a strong enrichment of Rop1-NeonFP at the parasite’s apical end. By comparison the apical:basal ratio in IAA treated parasites was only 2.1±0.15. While Rop1 is still enriched at the parasites apical end, there is an increase in Rop1-NeonFP fluorescence at the basal end of TgMyoF-KD parasites compared to controls (Fig. 9G).

**Figure 9.**
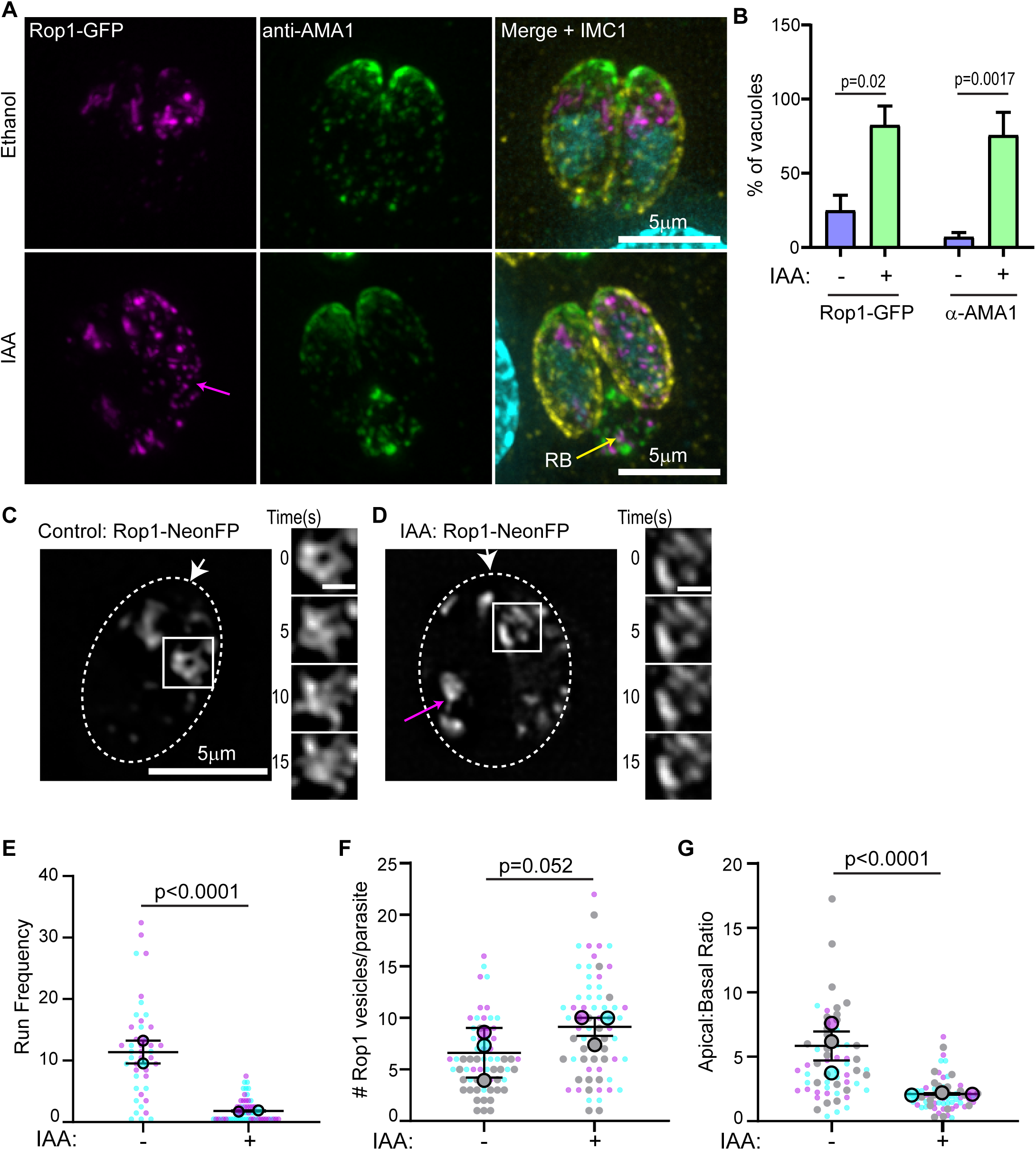
TgMyoF depletion affects apical positioning of the rhoptries and Rop1 vesicle movement. (A) TgMyoF-AID parasites expressing Rop1-NeonFP (magenta) were treated with EtOH and IAA for 15 hours before fixation and staining with an anti-AMA1 (green) and anti-GAP45 (yellow) antibodies and DAPI (cyan). Magenta arrow indicates Rop1 vesicles in the cytosol. Yellow arrow indicates the residual body (RB). (B) Quantification of TgMyoF-AID parasites with micronemes (visualized with anti-AMA1 antibody) and rhoptries (visualized by expression of Rop1-NeonFP) in the residual body. N = 57 vacuoles for AMA-1 quantitation, N=77 vacuoles for Rop-1 quantitation from two independent experiments. (C&D) TgMyoF-AID parasite’s expressing Rop1-NeonFP were treated with EtOH or IAA for 15 hours before live cell imaging. *Right panel:* Rop1 localization. Magenta arrow indicates rhoptry mislocalization to the parasites basal end. White arrow indicates parasite apical end. Dashed oval indicates the PV in a 2-parasite vacuole. Area in the white box was used to make inset in the left panel. *Left panel:* Dynamics of rhoptries in the first 15 seconds of imaging. (E) Run frequency (# of directed runs/parasite/minute) of Rop1 vesicles in control and TgMyoF depleted parasites. See table S1 for more details. (F) Number of Rop1 vesicles per parasite. N = 63 and 76 parasites in control and TgMyoF-KD parasites respectively in 3 independent experiments. (G) Ratio of Rop1 fluorescence in the apical and basal ends of the parasite. N = 63 and 76 parasites in control and TgMyoF-KD parasites respectively in 3 independent experiments. (E-G) Mean from each independent experiment is indicated with large circles. Raw data is shown with smaller circles; experiment 1 in magenta, experiment 2 in cyan, experiment 3 in grey. Error bars indicate mean and SEM.

## Discussion

The polarized endomembrane system in *T. gondii* is vitally important for the accurate trafficking of secretory proteins to the micronemes, rhoptries and dense granules. Our data demonstrate that F-actin and TgMyoF control the dynamics, positioning and morphology of the endomembrane network in *T. gondii*.

Actin-driven ER tubule motility observed in *T. gondii* is reminiscent of the microtubule-controlled ER tubule movements that have been described in mammalian cells (60,61,65) and actin/myosin XI driven tubule movements in plants (66). The function of ER tubule movement is best understood in mammalian cells, where ER tubules form contacts with numerous organelles and regulates the timing and site of mitochondrial and endosome fission, as well as facilitating lipid and calcium exchange between organelles (67–69). Future work is required to elucidate the importance of ER tubule movement in *T. gondii* biology.

The Golgi in *T. gondii* is localized adjacent to the nuclear envelope at the parasite’s apical end, appearing as a single stack when imaged by conventional fluorescence microscopy. Early in the parasite’s cell division cycle, the Golgi elongates and divides by binary fission (52,63). A second round of Golgi division then occurs, and two Golgi are inherited by each daughter (51) which appear to coalesce to form a single Golgi stack. However, it remains undetermined whether the two Golgi stacks undergo membrane fusion or if they are simply maintained in close proximity such that the stacks cannot be resolved using diffraction-limited microscopy. In this study, we quantified Golgi number four hours after TgMyoF depletion and one hour after actin depolymerization with cytochalasin D. Golgi fragmentation had occurred at each of these time points, with no accompanying effect on centrosome number suggesting the changes in Golgi number are not due to a perturbation of the parasite’s cell division cycle. One possible explanation for these results could be that the Golgi may be comprised of a paired stack held together in very close proximity in an actin-dependent manner. A similar observation was recently made by Kondylis and colleagues who demonstrated in Drosophila S2 cells that the Golgi consists of duplicated structural units that separated during the G2 phase of the cell cycle, prior to mitosis (70). Golgi separation could be artificially induced upon actin depolymerization with either cytochalasin D or latrunculin B (LatB). The observation that Golgi number did not increase between the 4- and 15-hour IAA treatment times suggests that the number of structural units that can be formed from a single Golgi is finite. In addition to the increase in Golgi number, many Golgi failed to maintain their position at the parasite’s apical end and were observed associated with the lateral of basal sides of the nucleus (Fig. 8B and 8E). Collectively, our data demonstrate that actin and TgMyoF control both Golgi number and positioning.

After exiting the Golgi, proteins destined for the micronemes and rhoptries are trafficked to one or more post-Golgi compartments. We have identified Rab6 as a new marker of the syntaxin 6 compartment, a protein that plays a role in retrograde trafficking from the Rab5a/Rab7 compartments to the Golgi (25). Our data builds upon previously published results indicating that Rab6 localizes to cytosolic vesicles and a compartment at the apical end of the nucleus, thought previously to be the Golgi since TgRab6 localizes to the Golgi when heterologously expressed in mammalian cells (25). However, we find no evidence that Rab6 is found in the cis-or mid-Golgi. Additionally, while we also observe Rab6(+) vesicles throughout the parasite, Rab6 does not colocalize with a marker for the dense granules, indicating these are distinct vesicle types. This data is consistent with a recent report demonstrating that Rab11a is found on the surface of dense granules and required for their secretion (71). The discrepancy between our results and previously published data, is likely due to the unavailability of organelle markers at the time of the previous publication (25).

The Rab6 compartments is dynamic and has tubular morphology that undergoes continuous rearrangement, we observed new tubules growing from the compartment while others retract. Vesicles were observed to bud from the tip of tubules and subsequently exhibited directed actin and TgMyoF dependent movement. This dynamic morphology closely mirrors the dynamics of endosomal compartments in mammalian cells. In this case, tubule formation is driven by the coordinated activity of microtubule motors (72) and branched actin networks nucleated by WASH and the Arp2/3 complex (73,74), along BAR domain-containing proteins that induce membrane curvature (75). No recognizable BAR-domain containing proteins or Arp2/3 complex proteins are found in *T. gondii*, (76,77) indicating that the molecular mechanisms underlying tubule formation in the Rab6 compartment are distinct from the mechanisms of tubule formation in mammalian cells and will require further investigation.

The apical localization of the Rab6/Syn6, Rab5a, Rab7 and DrpB compartments are all dependent on F-actin and TgMyoF. This dependence on TgMyoF and actin for apical positioning suggests that these post-Golgi compartments are associated with the actin cytoskeleton, or that the physical connections between these compartments, that maintain the compartments in close proximity at the apical end, are formed in an actin dependent manner. Future studies are needed to further identify the molecular players that control the associations between these compartments.

Formation of rhoptry organelles is cell cycle regulated and occurs in the mid to late stages of daughter cell development (31,52). Upon exit from the Golgi, rhoptry proteins are trafficked through the Rab5 compartment and then processed in a premature-rhoptry from which the mature rhoptry develops (10,22). Rhoptries had previously been shown to accumulate in the residual body upon actin depolymerization or TgMyoF knockdown, implicating TgMyoF in rhoptry trafficking and/or apical anchoring. Here, we further characterized the role of TgMyoF in rhoptry trafficking by imaging rhoptry dynamics in parasites expressing Rop1-NeonFP using live cell microscopy. As observed with the Rab6 compartment, the rhoptries undergo continual rearrangement and Rop1(+) vesicles were observed budding from the rhoptry bulb. Rop1 positive vesicles exhibited directed, TgMyoF-dependent movement throughout the parasite cytosol. In the absence of TgMyoF, there was a significant decrease in the number of directed runs exhibited by Rop1-NeonFP vesicles. Currently, we do not know the function or subcellular destination of these rhoptry derived vesicles and further work will be required to elucidate their biological role. These results suggest that although the rhoptries are formed once per cell cycle, acto-myosin dependent trafficking of proteins to or from the rhoptries occurs continuously.

There is an incomplete understanding of the mechanisms by which the mature rhoptries are anchored to the apical end of the parasite. In the absence of TgARO1, a membrane associated rhoptry protein, the apical positioning of the rhoptries was lost completely (78). This contrasts with the TgMyoF knockdown phenotypes where there is increased accumulation of rhoptries throughout the parasite cytosol, shown by an increase in Rop1 fluorescence at the parasite’s basal end (Fig. 9G), even though most parasites retain at least some intact rhoptries at the apical end. Thus, the TgARO1 knockdown parasites have a more severe rhoptry localization defect than TgMyoF knockdown parasites. Although TgMyoF was shown to interact indirectly with TgARO1 (78), the differences in the severity of these phenotypes suggest that TgMyoF is not required for TgARO1 anchoring activity. Our data suggests that TgMyoF is required for movement of immature rhoptries to the apical tip but is not required for TgARO1-dependent anchoring once the organelles have reached their destination.

This study demonstrates that TgMyoF controls the dynamics, positioning, and movement of a wide array of organelles in the endomembrane pathway in *T. gondii*. Future studies will be important to elucidate the mechanism by which this single molecular motor controls the movement of such a wide array of membranous cargos. Given the structural similarity between TgMyoF and the well characterized cargo transporter myosin V, we previously hypothesized that TgMyoF bound dense granules via its C-terminal WD40 domain and transported cargo by moving processively on filamentous actin (39). The large number of membrane-bound organelles whose movement is dependent on TgMyoF makes elucidating TgMyoFs mechanism of action even more pertinent. Outstanding questions include: Does TgMyoF associate directly with each membrane bound organelle? If so, what is the molecular basis of this association? How do these molecular complexes vary for each cargo and how are these interactions regulated? Does TgMyoF have the capacity to transport cargo as either a single motor or an ensemble? Future work aimed at identifying TgMyoF interacting proteins and modes of regulation will provide new insight into mechanisms of cargo transport in *T. gondii*.

## Materials and Methods

### Cell culture and parasite transfection

*T. gondii* tachyzoites derived from RH strain were used in all experiments. Parasites were maintained by continuous passage in human foreskin fibroblasts (HFFs) in Dulbecco’s Modified Eagle’s Media (DMEM) (ThermoFisher, Carlsbad CA) containing 1% (v/v) heat inactivated fetal bovine serum (FBS) (VWR, Radnor PA), 1X antibiotic/antimycotic (ThermoFisher) as previously described (79). Parasites were transfected as described previously (79) using a BTX electroporator set as follow: voltage 1500V; resistance 25Ω and capacitance 25μF.

### Drug treatment

To determine the effect of actin depolymerization on endomembrane organization and dynamics, transfected parasites were grown for 15-18 hours in confluent HFF monolayers, treated for 60 minutes with either 2µM cytochalasin D or equivalent volume of DMSO before live cell imaging as described below. To deplete TgMyoF, TgMyoF-AID parasites were treated with a final concentration of 500µM IAA, diluted 1:1000 from a 500mM stock made in 100% EtOH. For live cell imaging experiments treated time ranged from 15-18 hours. For western blot, treatment time was varied as indicated in figure 3.

### Construction of expression plasmids

A list of plasmids, primers and gene accession numbers used in this study can be found in Tables S2, S3 and Table S5 respectively.

#### Creation of pTKOII-MyoF-mAID-HA

pTKOII-MyoF-EmeraldGFP (EmGFP) (39) was digested with BglII and AflII to remove the EmGFP coding sequence. AID-HA was amplified by PCR using the AID-HA ultramer as a template and primer pairs AID-HA F and AID-HA R. Plasmid backbone and the PCR product was gel purified and ligated via Gibson assembly using NEBuilder HiFi DNA assembly master mix as per manufacturer’s instructions (New England BioLabs; Ipswich, MA). Plasmids were transfected into NEB5α bacteria and positive clones screened by PCR and verified by Sanger sequencing.

#### Creation of pmin-eGFP-Rab6-Ble

To create parental plasmid pmin-eGFP-mCherry-Ble, pmin-eGFP-mCherry was digested with KpnI and XbaI. pGra1-Ble-SAG1-3’UTR plasmid (80) was digested with KpnI and XhoI to remove the ble expression cassette. A fill-in reaction was performed to produce blunt ends by incubating plasmids with 100µM dNTPs and T4 DNA polymerase at 12°C for 15 minutes. Digested plasmids were gel purified and ligated together using T4 DNA ligase (New England Biolabs). Plasmids were transfected into NEB5α bacteria and positive clones screened by PCR and verified by Sanger sequencing. To create pmin-EmGFP-Rab6-ble, pmin-eGFP-mCherry-Ble was digested with NheI and AflII to remove eGFP-mCherry sequence. EmGFP was amplified by PCR with EmGFP-R6F and EmGFP-R6R primer pairs using pTKOII-MyoF-EmGFP as a template. Rab6 coding sequence was amplified by PCR using RH cDNA as a template and Rab6F and Rab6R primers. Digested plasmid backbone, EmGFP and Rab6 PCR products were gel purified and ligated using NEBuilder HiFi DNA assembly master mix as per manufacturer’s instructions (New England BioLabs). Plasmids were transfected into NEB5α bacteria and positive clones screened by PCR and verified by Sanger sequencing.

#### Creation of ptub-SAG1-ΔGPI-HDEL

Parental plasmid ptub-SAG1-ΔGPI (39) was digested with AflII. HDEL ultramers (Table S3) reconstituted to a concentration of 200mM in duplex buffer (100mM K Acetate; 30mM Hepes pH 7.5), were combined in equal ratios and headed to 95°C for 5 minutes before cooling slowly to room temperature. Duplexed ultramers were diluted 1:100 in molecular biology grade water and ligated to digested plasmid using NEBuilder HiFi DNA assembly master mix as per manufacturer’s instructions (New England BioLabs). Plasmids were transfected into NEB5α bacteria and verified by Sanger sequencing.

#### Creation of pmin-NeonFP-Rab5a and pmin-NeonFP-Rab7

Rab5a and Rab7 coding sequences were amplified by PCR using primers sets Rab5F/R, Rab7F/R and pTg-HARab5a and pTg-HARab7 (27,81) as templates. NeonGreen was amplified by PCR using Neon-R5F/Neon-R5R, Neon-R7F/Neon-R7R primer sets and Ty1-NeonGreenPave as a template. pmin-EmGFP-Rab6 plasmid was digested with NheI and AflII to remove EmGfP-Rab6 coding sequence. Plasmid backbones and PCR products were gel purified and annealed using Gibson assembly with NEBuilder HiFi DNA assembly master mix as per manufacturer’s instructions. Plasmids were transfected into NEB5α bacteria and positive clones screened by colony PCR and verified by Sanger sequencing.

#### Creation of ptub-Rop1-NeonGreenFP

ptub-SAG1-ΔGPI-GFP plasmid was digested with NheI and AflII to remove SAG1-GFP coding sequence. Rop1-GFP coding sequence was amplified using Rop1 F/R primer pairs and RH cDNA as a template. NeonGreen was amplified by PCR using NeonRop1F and NeonRop1R primer pairs. Plasmid backbones and PCR products were gel purified and annealed using Gibson assembly with NEBuilder HiFi DNA assembly master mix as per manufacturer’s instructions. Plasmids were transfected into NEB5α bacteria and positive clones screened by PCR and verified by Sanger sequencing.

### Creation of TgMyoF-AID parasite line

The *pTKO2_MyoF_mAID-HA* plasmid was linearized with SphI and 25μg was transfected into 1×10^7^ Δ*Ku80:ΔHXGPRT:Flag-Tir1* parental parasites (a gift from Dr. David Sibley, Washington University (56)). Parasites were selected using mycophenolic acid (MPA) (25 µg/ml) and xanthine (50 µg/ml) until approximately 70% of the parasites were HA positive. Clonal parasite lines were obtained by limited dilution into a 96 well plate. After 7 days of growth, wells containing a single plaque were selected for further analysis. All HA positive clones were amplified in a 6 well plate and genomic DNA isolated using Qiagen DNAeasy blood and tissue kit as per manufactures instructions (Qiagen, Germantown, MD) (Cat #69504). Genomic DNA was analyzed for correct insertion of the pTKO2_MyoF_mAID HA plasmid into the TgMyoF genomic locus by PCR using primers listed in Table S3 as outlined in Fig. S1.

### Western blot

To assess the extent of TgMyoF depletion after auxin addition as a function of time, 8×10^6^ extracellular TgMyoF-AID parasites were incubated in 500µl of DMEM containing 1% FBS, antibiotic/antimycotic and 500μM IAA and incubated at 37°C, 5% CO_2_ for 0, 0.5, 1, 2, or 4 hours. At each time point parasites were centrifuged at 1200xg for 5 minutes and resuspend in 25ul of 1xPBS containing protease inhibitor cocktail (MilliporeSigma, St. Louis MO; Cat # P8340). 25ul of 2XLamelli buffer (Biorad, Hercules, CA; Cat # 1610737) containing 100mM DTT was added to each sample and boiled at 95°C for 10 minutes. Samples were run on 4-12% gradient gel (Biorad; Cat# 456-1064), transferred to PVDF for 60 minutes at 4°C at 100V using the Biorad blotting system. PVDF membranes were blocked overnight at 4°C in 5% milk, 1XTBS-T (150mM NaCl, 20mM Tris base, 0.1% tween20 (v/v) pH 7.4) before blotting with tubulin and HA primary antibodies and anti-mouse and anti-rat secondary antibodies. All antibodies were incubated with PVDF for 60 minutes at room temperature with 3×10-minute washes with 1xTBS-T in between antibody incubations. Immunoblots were developed using Pierce ECL western blotting substrate (ThermoFisher, Cat# 32209) and visualized using LiCor Odyssey Fc. Antibody dilutions and product information can be found in Table S4.

### Microscopy

#### Parasite transfection for live cell imaging or immunocytochemistry

25µg of each plasmid was transfected as described above. Transfected parasites were grown for 15-18 hours in confluent HFF monolayers grown on either MatTek dishes (MatTek corporation, Ashland MA) or on coverslips before either live cell imaging or immunocytochemistry.

#### Live cell microscopy

Growth media was replaced with Fluorobrite DMEM (ThermoFisher; Cat# A19867) containing 1% FBS and 1x antimycotic/antibiotic pre-warmed to 37°C. Images were acquired on a GE Healthcare DeltaVision Elite microscope system built on an Olympus base with a 100x 1.39 NA objective in an environmental chamber heated to 37°C. This system is equipped with a scientific CMOS camera and DV Insight solid state illumination module. Image acquisition speeds were determined on a case-by-case basis as noted in the video legends.

#### Immunocytochemistry

Parasites were fixed with freshly made 4% paraformaldehyde (Electron microscopy sciences, Hatfield, PA; Cat# 15714) in 1xPBS (ThermoFisher; Cat# 18912-014) for 15 minutes at RT. Cells were washed three times in 1xPBS and permeabilized in 0.25% TX-100 diluted in 1xPBS for 10 minutes at room temperature before washing three times in 1xPBS. Cells were blocked in 2% BSA-1XPBS for 15 minutes before antibody incubations. All antibodies were diluted in 0.5% BSA-1xPBS at the concentrations indicated in Table S4. DNA was stained with 10μM DAPI diluted in 1xPBS for 10 minutes and then washed three times in 1xPBS. Cells in mattek dishes were either imaged immediately or stored in 1xPBS at 4°C. Coverslips were mounted onto slides using Prolong Gold anti-fade reagent (ThermoFisher; Cat # P36930) and allowed to dry overnight before imaging.

### Image analysis

#### Vesicle tracking and counting

Vesicle tracking was performed using MtrackJ plug-in in Fiji (National Institutes of Health) as previously described (82). Line scan analysis was performed using the “plot profile” tool in Fiji. Kymographs were made using Fiji plugin KymographBuilder. Fiji plugin cell counter was used to quantify the number of vesicles/parasites. Statistical significance was determined using students t-test.

#### Effect of TgMyoF depletion on post-Golgi compartment number

To count number of PGC “objects” in each parasite in control and TgMyoF knockdown parasites, transfected parasites were fixed and stained with the anti-IMC1 antibody and DAPI and z-stack images were acquired using Deltavision elite imaging system. For each PGC image, maximum intensity projection of Z-stacked images were converted to binary images using Fiji and particles counted using the “analyze particles” tool. IMC1 staining was used to determine number of parasites/vacuole. Statistical significance was determined using students t-test.

#### Quantification of Golgi and centrosome number

To quantify the number of Golgi and centrosome per parasite, TgMyoF-AID parasites were transfected with Grasp55-GFP plasmid alone or Grasp55-mCherry and Centrin1-GFP plasmids, grown in confluent HFF monolayers overnight and treated with either EtOH or IAA for the final 4- or 15-hours of growth before fixation and processing for immunofluorescence. Number of Golgi per parasite and number of centrosomes per parasites were counted manually using the cell count tool in Fiji. Statistical significance was determined using students t-test.

#### Quantification of rhoptry and micronemes in the residual body

To quantify the number of parasites with microneme and rhoptry proteins in the residual body, TgMyoF parasites were transfected with Rop1-NeonFP plasmid, and were grown for 15 hours in either EtOH or IAA before fixation and immunocytochemistry with an anti-AMA1 antibody (Table S4). The number of vacuoles containing rhoptries or micronemes in the residual body were manually counted. N = 57 vacuoles for AMA-1 quantitation, N=77 vacuoles for Rop-1 quantitation from two independent experiments. Statistical significance was determined using students t-test.

#### Loss of rhoptry positioning at the parasites’ apical end

To quantify Rop1 localization at the apical or basal ends of the parasite, Rop1-NeonFP was transiently expressed in TgMyoF-AID parasites treated with EtOH or IAA for 15 hours. Parasites were imaged live as described above. Using the first frame of each movie, the apical half and the basal half of each parasite was outlined manually and the mean fluorescent intensity of Rop1 in the apical and basal ends of the parasites were calculated using Fiji. The apical to basal mean fluorescence intensity was calculated for each parasite. A ratio of 1 indicates that Rop1 is evenly distributed in the apical and basal ends. A ratio >1 indicates more rop1 is localized in the apical end than the basal end and a ratio <1 indicates more rop1 in the basal end compared to the apical end. Rop1-NeonFP fluorescence in the residual body was excluded from this analysis. Statistical significance was determined using students t-test.

### Statistics

Statistical analyses were performed using GraphPad Prism. Superplots were made as described in (83).

## Funding Information

This work was funded by National Institutes of Health R21AI121885 awarded to ATH and the University of Connecticut Research excellence program awarded to ATH. The funders had no role in study design, data collection and interpretation, the decision to submit the work for publication or manuscript preparation. The authors declare that no competing interests exist.

## Acknowledgements

We thank members of the Heaslip Lab, Dr. Ken Campellone and members of the Campellone lab (University of Connecticut) for helpful discussion during the course of these experiments. We thank our colleagues for sharing reagents: Dr. Gary Ward (University of Vermont) for the IMC-1 and AMA-1 antibodies; Dr. Markus Meissner (Ludwig-Maximilian University, Munich) for the DrpB and Syntaxin 6 expression constructs; Dr. Vern Carruthers (University of Michigan) for the TgCPL antibody and the Rab5 an Rab7 expression plasmids. Dr. David Sibley (Washington University, St. Louis) for sharing the Flag-Tir1 parasite line; Dr. Boris Striepen (University of Pennsylvania) for the anti-Cpn60 antibody.

## Notes

### Competing Interest Statement

The authors have declared no competing interest.

### Summary of Updates

Version 1 was transferred directly from journal website and figure images were not transferred. Version 2 contains all main text figures.

